# Why is the explicit component of motor adaptation limited in elderly adults?

**DOI:** 10.1101/753160

**Authors:** Koenraad Vandevoorde, Jean-Jacques Orban de Xivry

## Abstract

The cognitive component of motor adaptation declines with aging. Yet, in other motor tasks, older adults appear to rely on cognition to improve their motor performance. It is unknown why older adults are not able to do so in motor adaptation. In order to solve this apparent contradiction, we tested the possibility that older adults require more cognitive resources in unperturbed reaching compared to younger adults, which leaves fewer resources available for the cognitive aspect of motor adaptation. Two cognitive-motor dual-task experiments were designed to test this. The cognitive load of unperturbed reaching was assessed via dual-task costs during the baseline period of visuomotor rotation experiments, which provided us with an estimation of the amount of cognitive resources used during unperturbed reaching. However, since we did not observe a link between dual-task costs and explicit adaptation in both experiments, we failed to confirm this hypothesis. Instead, we observed that explicit adaptation was mainly associated with visuospatial working memory capacity. This suggests that visuospatial working memory of an individual might be linked to the extent of explicit adaptation for young and older adults.

## Introduction

Several characteristics of movement are affected by aging, such as movement speed (Diggles-Buckles, 1993), reaction times (Fozard et al., 1994), and motor coordination (Serrien et al., 2000). These movement changes result from a broad range of physiological (e.g. muscle fatigue, muscle atrophy, sensory function decline, cartilage reduction) (Jubrias et al., 1997; Lindle et al., 1997; Martin and Buckwalter, 2002; Goble et al., 2009; Owsley, 2011) and brain changes (e.g. brain atrophy, altered brain connectivity) (Guttmann et al., 1998; Raz et al., 2005; Davis et al., 2012). Age-related movement deficits are already present during simple point-to-point reaching movements, occurring as increased spatial and temporal movement variability and movement slowing (Darling et al., 1989; Yan et al., 1998; Ketcham et al., 2002; Van Halewyck et al., 2015). Apart from the movement parameters that are altered with aging, controlling movement might also impose increased cognitive demand on older adults compared to younger adults. This might be the result of sensory acuity and body physiology changes with aging, likely making the same task more challenging for elderly (Li and Lindenberger, 2002; López-Otín et al., 2013). Increased cognitive demand is also reflected by increased neural recruitment with aging (Heuninckx et al., 2005, 2008). This age-related over activation has been described to be compensatory in order to obtain better task performance (Reuter-Lorenz and Campbell, 2008). In addition, available capacity of cognitive resources is most likely lower in elderly adults because prefrontal cortex degrades with aging and through age-related brain atrophy, altered brain connectivity, and changes in corpus callosum (Ota et al., 2006; Seidler et al., 2010). In motor tasks, older adults are thought to rely more on cognition to improve their motor performance (motor coordination: (Heuninckx et al., 2008); Balance:(Boisgontier et al., 2013)). This suggests that elderly adults might also use more cognitive resources for simple reaching movements.

Besides altered characteristics of movement, motor learning is also affected by aging. One specific form of motor learning is motor adaptation. Motor adaptation is the adjustment of movement to changes in the environment or to body changes (Shadmehr et al., 2010). Motor adaptation can be divided in multiple components; such as the explicit and implicit components of motor adaptation (Mazzoni and Krakauer, 2006; Benson et al., 2011; Taylor et al., 2014; Mcdougle et al., 2016). The explicit component of adaptation is the part of adaptation that is under cognitive control, while the implicit component is the adaptation that takes place without conscious awareness of the participant. Several ways exist to measure these two components: Taylor et al. (2014) asked participants to report their aiming direction, Morehead et al. (2015) and Werner et al. (2015) asked participants to switch on or off their aiming direction together with a switch in color cue, and Morehead et al. (2017) introduced task-irrelevant cursor feedback that caused a drift in hand movement away from the visual error. Motor adaptation declines with aging and this decline can be attributed to the decline of the explicit component of motor adaptation (Heuer and Hegele, 2008; Vandevoorde and Orban de Xivry, 2019). A higher capacity of working memory is linked to a larger explicit component of motor adaptation, while differences in working memory capacity are not linked to the implicit component (Christou et al., 2016). Explicit strategies are useful to speed up the learning process since the explicit learning rate is higher than the implicit learning rate (McDougle et al., 2015). However, this faster learning in motor adaptation can be impaired by introducing a second cognitive task (Taylor and Thoroughman, 2008; Galea et al., 2010; Keisler and Shadmehr, 2010; Malone and Bastian, 2010), and is likely to occur through disturbance of the explicit component.

Since elderly people rely more on cognition in motor tasks (Heuninckx et al., 2008; Boisgontier et al., 2013), it is unknown why older adults are unable to do so in motor adaptation, especially as previous evidence indicates that the cognitive part (i.e. the explicit component) of the motor task is affected. One possible explanation is that the increased reliance on cognition for simple reaching movements in elderly people would leave fewer cognitive resources available for concurrent tasks (here motor adaptation) as cognitive resources are limited (Li and Lindenberger, 2002; Heuninckx et al., 2005, 2008; Seidler et al., 2010). In order to investigate this explanation, we propose the ***cognitive resources hypothesis*** (Figure 1). The hypothesis suggests that older adults already require most of their available cognitive resources for simple reaching movements. The increased cognitive demand in “simple” reaching movements might in turn result in an age-related decline of explicit adaptation. This hypothesis is in line with the compensation-related utilization of neural circuits hypothesis (CRUNCH) that describes over- and under-recruitment of neural resources depending on the cognitive load in elderly (Figure 1). During a task with low levels of cognitive load, older adults tend to recruit more cognitive resources compared to younger adults to maintain task performance (Reuter-Lorenz and Campbell, 2008). Instead, during tasks with higher levels of loads, older adults cannot compensate their performance anymore and they recruit similar or fewer cognitive resources than younger adults (Grady, 2012).

**Figure 1:**
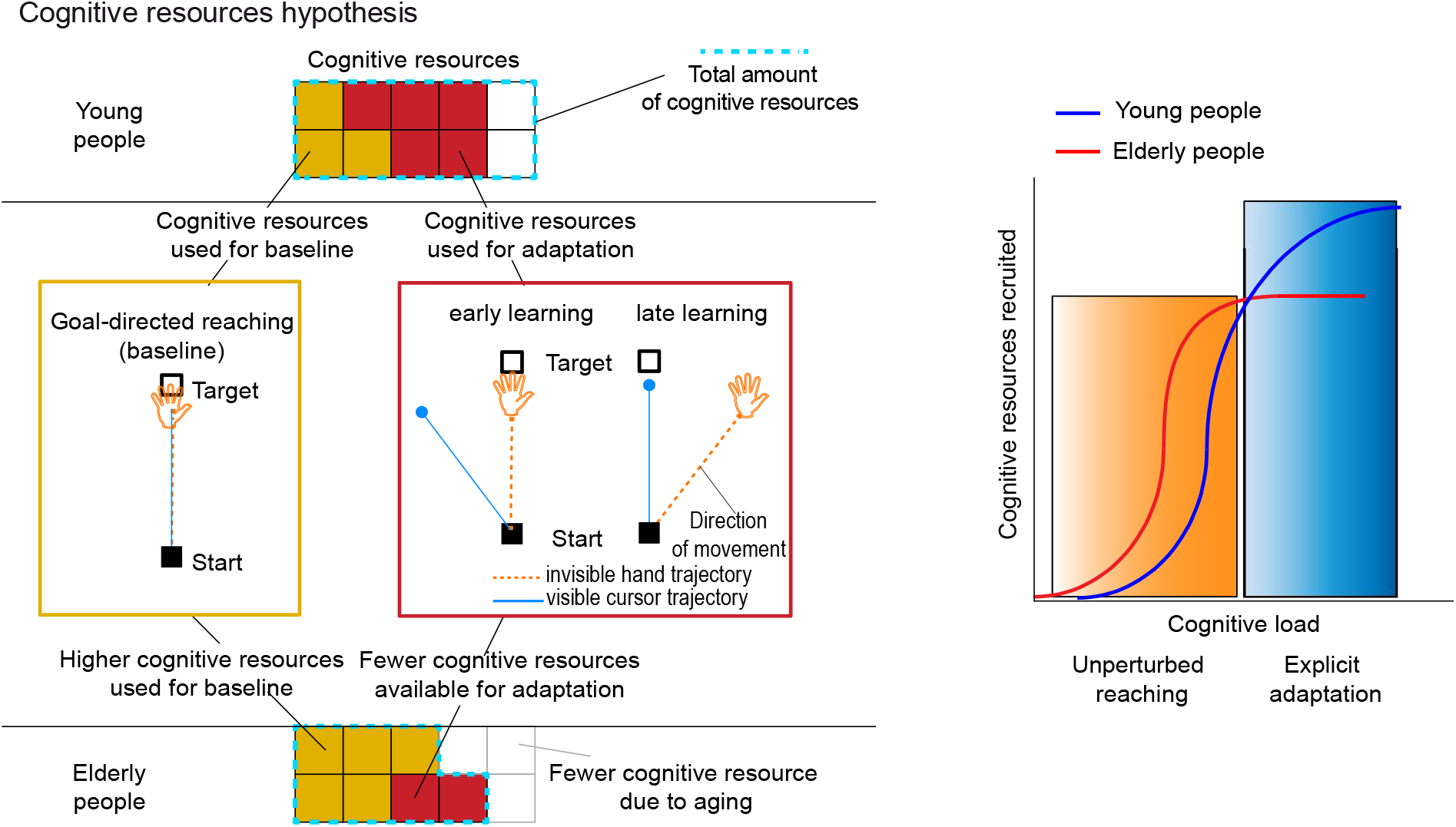
Cognitive resources hypothesis. Left part: **Cognitive resources hypothesis**: Elderly adults require more cognitive resources to perform goal-directed reaching compared to younger adults (orange blocks). The total amount of resources is reduced with aging (blue dotted area). As a result, fewer resources are available for explicit adaptation in the elderly. The elderly have a saturation of their resources when adjusting their movement (two red blocks). Young adults, instead, do not reach their total amount of available resources (five red blocks, still two additional blocks remain available). Right part: “compensation-related utilization of neural circuits hypothesis (CRUNCH)” (Figure adapted from *Grady, 2012*) applied to motor adaptation: In unperturbed reaching (i.e. lower cognitive load task) elderly people recruit more cognitive resources (orange box), while young people can apply more cognitive resources (blue box) during development of explicit adaptation (i.e. higher cognitive load task).

The increased (compensatory) brain activity in elderly is often located in frontal brain regions such as prefrontal cortex (Mattay et al., 2006; Schneider-Garces et al., 2009; Cappell et al., 2010) which is a region that is highly affected by aging as evidenced by structural (Raz and Rodrigue, 2006) and connectivity deficits (Madden et al., 2010; Nagel et al., 2011). At the same time, brain regions responsible for the explicit component of adaptation, are most likely frontal lobe regions, including lateral and medial aspects of prefrontal cortex as well as premotor cortex (Shadmehr and Holcomb, 1997; Krakauer et al., 2004; Mcdougle et al., 2016). Together, prefrontal cortex appears to be the region where both age-related compensatory brain activity takes place, and the region that might be responsible for the explicit component of adaptation. This might indicate that the same (prefrontal) cognitive resources are required for compensation with aging and are responsible for motor adaptation decline.

The amount of cognitive resources applied during a motor task can be assessed with cognitive-motor dual-task experiments (Boisgontier et al., 2013). In cognitive-motor dual-task experiments, a cognitive task is performed during a motor task which reduces motor and/or cognitive performance. The reduction of performance can be quantified as a dual-task cost (Li and Lindenberger, 2002; Boisgontier et al., 2013). Dual-task costs show that cognition is involved in motor tasks; this is the case even in natural motor tasks such as standing with stable posture (Boisgontier et al., 2013). A link between cognitive resources in baseline movement and explicit adaptation would show that the deficit of explicit adaptation with aging could be defined as a resource competition problem (Figure 1): Resources that are necessary for the adaptation process are already used for allowing simple reaching movement.

In order to answer this question, we designed two cognitive-motor dual-task experiments that allowed us to quantify the amount of cognitive resources used during simple reaching movements with a dual-task in the baseline of a motor adaptation experiment. A response inhibition task was selected as a cognitive task during the first dual-task experiment since it would reduce attention for the reaching task. This task requires responding quickly to visual stimuli, which could interfere with the visual feedback from the reaching task. At the same time, this task requires filtering distracting stimuli and is therefore suited to assess selective attention (Kopp et al., 1994). A working memory capacity task was selected as a cognitive task during the second dual-task experiment since working memory is known to be linked to explicit adaptation (Anguera et al., 2010, 2012; Christou et al., 2016). Therefore, the working memory task would interfere with resources that are required for motor adaptation.

## Material and methods

### Participants

A total of 143 healthy adults were recruited and participated after providing written informed consent. Eighty-one participated in experiment E1 and 62 participated in experiment E2.

All 81 participants were included in the final analyses for experiment E1. These 81 participants consisted of 41 young adults (between 20 and 35 years old, age: 23.1 ± 3.5 years, mean ± SD; 25 females) and 40 older adults (between 60 and 75 years old, age: 67.5 ± 4.5 years; 23 females). Sixty-two of the 81 included participants were the same participants as in paradigm 3 of Vandevoorde and Orban de Xivry (2019b). They participated in experiment E1b (same as cued visuomotor rotation in this paper) and E4 of this last-mentioned paper. Afterwards eleven young and ten older participants were subsequently recruited for the current study in the context of a master student project.

All 62 participants were included in the final analyses for experiment E2. These 62 participants consisted of 30 young adults (between 20 and 32 years old, age: 22.9 ± 2.7 years, mean ± SD; 15 females) and 32 older adults (between 61 and 75 years old, age: 67.6 ± 4.5 years; 13 females).

The Edinburgh handedness questionnaire (Oldfield, 1971) was used to confirm that all participants were right-handed. All participants were screened with a general health and consumption habits questionnaire. None of them reported a history of neurological disease or were taking psychoactive medication. In older adults general cognitive functions was assessed using the Mini-Mental State Examination (MMSE) (Folstein et al., 1975). All elderly scored within normal limits (MMSE-score ≥ 26). The protocol was approved by the local ethical committee of KU Leuven, Belgium (project number: S58084). Participants were financially compensated for participation (10 €/h).

### Experimental setup

Participants were seated in front of a table. With their right hand, participants performed a reaching task on a digitizing tablet (Intuos pro 4; Wacom) with a digitizing pen. A wooden cover above the tablet prevented participants from receiving visual feedback of their moving arm. Movement trajectories were recorded at 144 Hz. The only visual feedback was displayed on a 27 inch, 2560 x 1440 optimal pixel resolution LCD monitor with 144 Hz refresh rate (S2716DG, Dell), vertically mounted in front of the participant.

#### Assessment of dual-task cost

##### Overall dual-task design

In the flanker dual-task experiment (E1) (Figure 2A), the first part of the experiment (260 trials) was designed for assessing the dual-task cost during baseline of the cued motor adaptation task. Participants first performed 20 target reaching trials. Then, a block of 60 trials was repeated four times during the dual-task baseline (Figure 2A). The two separate tasks used in the dual-task were reaching to a target with the right (dominant) hand and a flanker task with the left hand. Each block of 60 trials consisted of the single reaching motor-task (20 target reaching trials), the single flanker cognitive-task (20 reaction time inhibition trials) and the cognitive-motor dual-task (20 combination of reaching and reaction time trials). Before the experiment, each participant performed 30 familiarization trials to make sure that they understood the instructions and that they performed the task correctly. These familiarization trials consisted of 10 target reaching trials, 10 flanker trials and 10 combination trials. A break of 60 seconds was applied before trial 141 of the dual-task.

**Figure 2:**
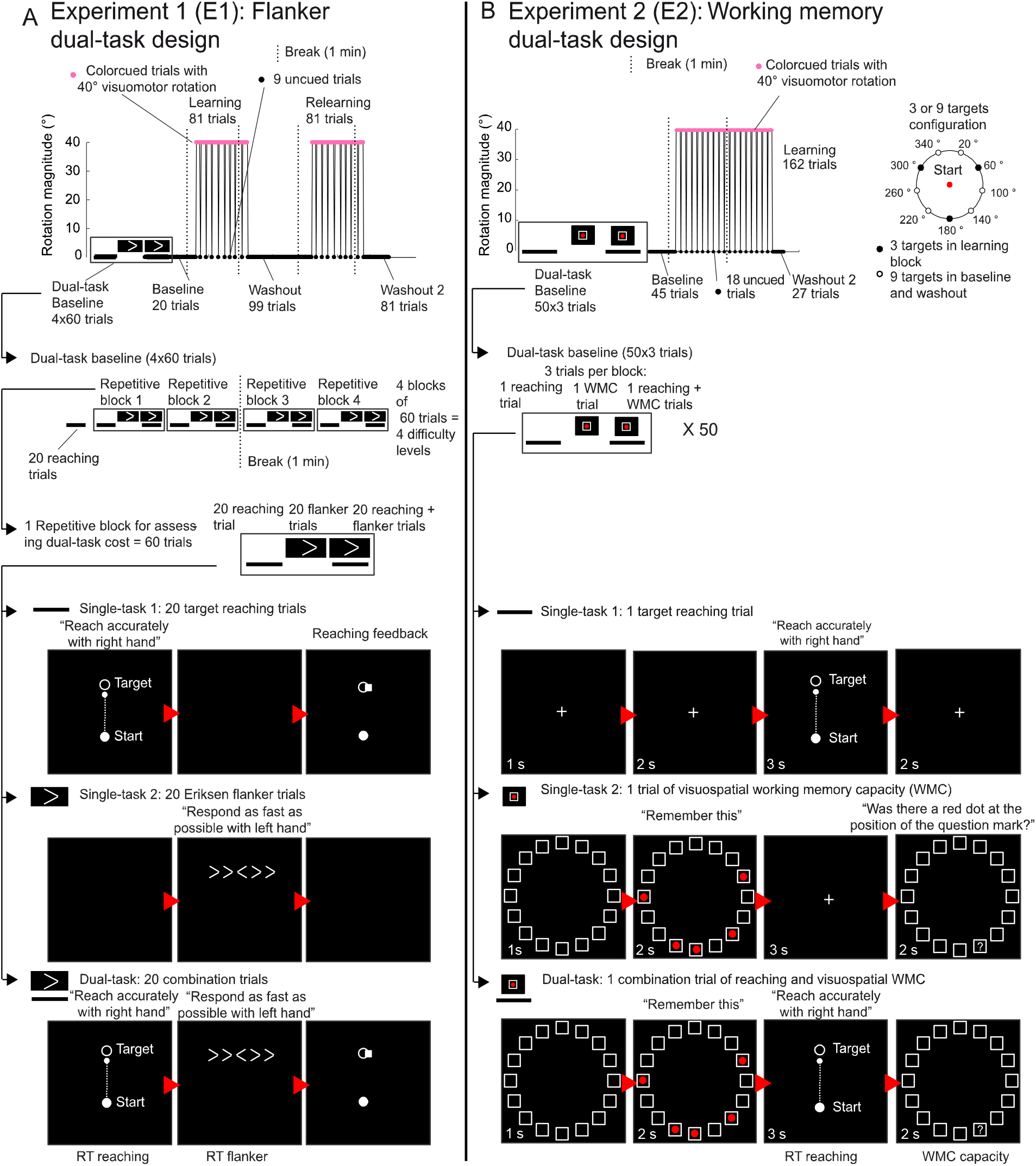
Cognitive-motor dual task paradigms. Explicit adaptation level was assessed with cued motor adaptation. A change in cursor color indicated the presence or absence of a 40 ° visuomotor rotation. In the baseline of the cued motor adaptation experiment, a dual-task was introduced to quantify the amount of cognitive resources applied during unperturbed reaching. Two designs of the dual-task were used: **A) Flanker dual-task design (E1).** The cognitive-motor dual-task consisted of a combination of an unperturbed target reaching task and a flanker task. During the flanker task, participants had to indicate the direction of the middle arrow among several arrows as fast as possible. **B) Working memory dual-task design (E2).** The cognitive-motor dual-task consisted of a combination of unperturbed target reaching and a visual working memory task. In the visual working memory task, participants had to remember positions of red dots, which were presented in a circular array. Afterwards they had to indicate whether a probed position contained a red dot. (1 s = 1 second)

Similar as in the flanker dual-task (E1), the first part of the working memory dual-task (E2) (Figure 2B) (150 trials) was designed for assessing the dual-task cost during baseline of the cued motor adaptation task. The two separate tasks used in the dual-task were target reaching and a working memory capacity task, both executed with the right dominant hand. The dual-task consisted of a single reaching motor-task, a single working memory capacity (WMC) cognitive-task and the two tasks combined. The cognitive-motor dual-task consisted of 150 trials, which was a repetition of 50 times a three-trial block (one target reaching trial, one working memory trial and one combination trial). At the start of the experiment, participants could practice the dual-task with five three-trial blocks, with exactly the same configuration as the real dual-task. This was important to get familiarized with every aspect: e.g. timing of task, answering for working memory capacity, reaching speed. Breaks of 30 seconds were introduced before trial 51, 102 of the dual-task.

##### Target reaching task (single motor-task)

For the target reaching, participants were instructed to make fast and straight shooting movements through the targets without stopping in the target. The participants held a digitizing pen in their right hand as if they were writing. They were instructed to always touch the surface of the tablet with the tip of this pen and to move their right arm and not only their wrist. The goal of each reaching movement was to slide through a target with a white cursor. Targets were presented at nine possible locations in E1 and ten locations in E2. The targets were evenly distributed on a circle 10 cm away from the starting position.

The reaching target was always a single white circular target. The diameters of the starting point and the target were both 10 mm and the feedback cursor had a diameter of 5 mm. The feedback cursor, which represented hand position was visible until movement amplitude exceeded 10 cm. At this point, a white square marked the position where movement amplitude reached 10 cm, providing visual feedback about the end point accuracy of the reach. The white square had sides of 5 mm and remained visible for 1.5 s. The zone for receiving 25 points was an additional 6 mm at both sides of the target. In the flanker dual-task (E1), while returning to the central starting position, the cursor disappeared and only a white arc (i.e. half circle) was visible. The radius of the return arc depended on the position of the pen on the tablet, i.e. the radius of the arc was equal to the radial distance between the position of the hand and the starting point. The center of the return arc was the central starting position. The reach area was divided in three different zones of 120 °. The arc was in the same 120° zone as where the participant’s (invisible) hand was. Participants had to move their hand in the opposite direction of the arc in order to return to the starting location. The arc allowed participants to return to the starting position and at the same time prevented the participants from using the visual feedback during the return movement to learn about the perturbation. In the working memory dual-task (E2), no return arc, but normal cursor feedback was used. To receive points, participants were required to reach the target between 175 and 375 ms after movement onset. If the reaching movement was too slow, a low pitch sound was played and the target color switched from white to purple. If the reaching was too fast, a high pitch sound was played and the target color switched from white to red.

In the working memory dual-task (E2), the only differences with the target reaching of the flanker dual-task design (E1) were the number of targets, the score system, and the timing of the trials. In this reaching task, in E2 ten targets were used instead of nine in E1. In E2, 50 single reaching trials were used, such that each target could be repeated five times. When hitting the target with the correct speed (i.e. between 125 and 375 ms), participants received 50 points. In E1, they could receive bonus points: When hitting targets at successive trials, they received 10 extra bonus points for each extra target-hit. In E2, they could not obtain bonus points. When reaching in close proximity of the target, they received 25 points and no bonus points. The cumulative score of all previous trials was displayed throughout the experiment. In flanker dual-task (E1), at the end of a target reach, the feedback cursor froze for 1.5 s in order to mark the reaching accuracy. In the flanker dual-task (E1), at the end of the feedback period, the participant had to move the tip of the pen back to center of the tablet and wait there between 350ms and 850ms (in steps of 50ms) in order to initiate the next trial. If participants were slower than 5 s, the next trial would automatically initiate. The timing of each reaching trial was strictly controlled in E2, such that one trial took exactly eight seconds while in E1 the duration of each trial could be different. In E2, a fixation cross was first displayed for three seconds. Then the participants had to reach before fixated again two seconds. Immediately after the reaching movement, extra fixation time was added in order to obtain a fixed time duration of eight seconds per trial (Figure 2B). In case, they exceeded the eight seconds per trial, the next trial would be initiated.

##### Cognitive task (single cognitive-task)

In E1, the cognitive task was a flanker task, adapted from Eriksen and Eriksen (1974). In this task, an uneven number of left or right pointing arrows were presented to the participant. Participants needed to answer with a left or right key press whether the middle arrow was pointing to the left or right direction. The arrows surrounding the middle arrow could either have congruent directions (>>>>>) or incongruent directions (>><>>) with respect to the middle arrow. Reaction times are reported to be lower with incongruent flanker arrows (Kopp et al., 1994). This task was executed with the left hand. The goal for the participant was to indicate the direction of the middle arrow as fast as possible. The next trial was initiated immediately after pressing the left or right key. All blocks consisted of 20 flanker trials with the middle arrow randomly presented to the left or the right side. In addition, each of these blocks contained 10 congruent and 10 incongruent trials. We repeated the same order (number of arrows and congruency) of 20 flanker trials for the single-task block and the subsequent dual-task block. The flanker task was modulated from the beginning towards the end of the dual-task baseline by gradually increasing the number of flanker arrows from block to block from two (first repetition) to eight (last repetition).

In E2, the cognitive task was a working memory capacity task (McNab and Klingberg, 2008; Christou et al., 2016). Sixteen (empty) white squares (1.9 cm x 1.9 cm) were presented in a circular array (11.2 cm diameter) for 1 second (Figure 2B) on a monitor. Two, three, four, five or six red circles (0.8 cm diameter) were visualized for two seconds in the 16 white squares with each red circle presented randomly in one of the 16 squares. Participants were asked to remember the positions of the presented red circles. After these two seconds, participants fixated on a white cross (0.6 cm x 0.6 cm) for three seconds. Afterwards they were asked whether a probed location corresponded to a position that contained a red circle before. They had two seconds to give their answer by moving a cursor to the right side, to the “yes answer”-target, or to the left side, which was the “no answer”-target. Participant could move the cursor on the monitor by moving the pencil on the tablet. In total, one trial had a fixed time duration of eight seconds. In total 50 trials were used for the single-task condition. Each trial contained two, three, four, five or six red circles (10 trials/condition) with all conditions randomly mixed. Of the 10 trials for each condition, five trials were “no answers” and five trials were “yes answers”.

##### Cognitive-motor dual-task design

In E1, a cognitive-motor dual-task trial was a combination of the above-described target reaching (motor) and flanker (cognitive) task in one trial. The trial started as a target reaching trial. However, at the end of the reach, arrows were presented at the position of the target, instead of the feedback on the reaching accuracy. This feedback was presented after the participants had indicated the direction of the middle arrow. Participants were instructed to first reach to the target as accurately as possible by moving the right hand and immediately afterwards perform the correct left/right key press as fast as possible with the left hand. The cognitive-motor dual-task was expected to have a negative effect on both motor and cognitive performance compared to single-task performance: a decreased accuracy or increased reaction time for the target reaching and a decreased accuracy or an increased reaction time for the flanker task. The reward-feedback (score system) was the same as in the single-task conditions. Since reward was implemented for the reaching trials but not for the flanker task, task priority might be expected in favor of the reaching task. The instructions did not specify task priority, but we asked participants to do their best for both tasks.

In E2, a cognitive-motor dual-task trial was a combination of the above-described target reaching (motor) and working memory (cognitive) task in one trial. A dual-task trial started with the presentation of 16 white squares for 1 second. Next, the two to six red dots were visualized for two seconds. Immediately after this, participants performed a target reaching movement, after which extra fixation time was added to obtain three seconds in total for the reaching part. Finally, they gave their answer to the working memory capacity task by moving the cursor to the “yes” or “no”-location. They had two seconds to execute this last part. Again, every trial had a strictly controlled time duration of eight seconds. We matched trial duration for the single- and dual-task condition in order to avoid that a difference in trial duration would cause a difference in performance between single- and dual-task trials. Participants performed 50 dual-task trials in total, such that every condition (2-6 red dots) of the working memory capacity trial could be repeated 10 times and each of the ten targets could be reached to five times. All conditions were pseudo-randomly mixed. The reward-feedback (score system) was the same as in the single-task conditions. In other words, the points obtained for one successful working memory trial was twice that for the points for one successful reaching trial. However, bonus points could also be obtained in the reaching trials. The instructions did not specify task priority. Participants were instructed to obtain as many points as possible by doing their best for both tasks.

#### Assessment of explicit and implicit component of adaptation with cued adaptation experiment

In E1, explicit adaptation level was assessed with cued motor adaptation. A change in cursor color indicated the presence or absence of a 40 ° visuomotor rotation. The cued motor adaptation paradigm consisted of a baseline block of 20 trials, a learning block of 81 trials (9 cycles), a first washout of 99 trials (11 cycles), a relearning block of 81 trials (9 cycles) and a second washout of 81 trials (9 cycles). A single start point location (filled red circle in Figure 2B) was used. Nine targets were presented during reaching trials of the dual-task baseline and during baseline and washout trials of the cued motor adaptation paradigm. Three targets (filled black circles in Figure 2B) were used during learning blocks of the cued motor adaptation paradigm.

In E2, immediately after the dual-task baseline, a short reaching baseline of 45 trials (5 cycles) was implemented. A learning block of 162 trials (18 cycles) and a short washout of 27 trials (3 cycles) followed the baseline. The learning block was designed to assess explicit adaptation by using the cued motor adaptation paradigm. Target configuration for cued motor adaptation was the same as in E1 (Figure 2B).

Cued motor adaptation was designed for assessing implicit and explicit adaptation, the methods and results of this cued motor adaptation experiment were already described in Vandevoorde and Orban de Xivry (2019b) as experiment E1b. It is a visuomotor rotation experiment, adapted from experiment 4 of (Morehead et al., 2015) with a perturbation magnitude of 40 °.

During baseline and washout blocks of the cued adaptation task of experiment E1, we used the same nine targets as in target reaching, spaced 40° apart (from 20° to 340°). The targets were presented pseudo-randomly in cycles during baseline and washout with each of the nine targets presented once per cycle. During learning blocks, only three targets were used, spaced 120° (60°,180°,300°) (filled black targets). Therefore, in the learning blocks, 9-trial-cycles consisted of three 3-trial-subcycles. In each subcycle each of the three targets was presented once. Both learning blocks consisted of 9 cycles (or 27 subcycles or 81 trials). The cursor dot remained white during the baseline and washout blocks. In contrast, during the two adaptation blocks, the cursor became a pink square (i.e. cued trial) instead of a white cursor dot. This cue indicated the presence of a 40° rotation. In each adaptation block, the cursor became again a white cursor dot (i.e. uncued trials) for a few trials, indicating the absence of the perturbation. The instructions were: “First, the cursor will be a white dot, but at some point the cursor will change to a pink square. At that moment, something special will happen but you still have to try to do the same thing, reach to the target with the cursor. The cursor will sometimes change back to a white dot. These trials with a white dot are normal reaching trials like in baseline.” The change in behavior induced by the cue was thus a measure of the explicit component of adaptation as participants could use the cue to switch off any conscious strategies they were applying to counteract the perturbation (Morehead et al., 2015). We reinforced the awareness of cue switches (signaling a cued trial among uncued ones or an uncued trial among cued ones) with a warning sound played for each cue switch and with a text that indicated the cue switch, displayed for 5 s: ‘Attention! The color of the cursor has changed.’ Nine uncued trial were presented per adaptation block (trials 7, 16, 25, 35, 45, 53, 61, 72 and 81). These uncued trials were equally distributed among the three targets (three uncued trials per target). In E1, breaks of 60 seconds were introduced in the cued adaptation experiment before trial 15, 95, 175 and 275. These breaks were respectively 5 trials before the onset of the first perturbation block, 6 trials before onset of the first washout block, 25 trials before the onset of the second perturbation and 6 trials before the onset of the second washout.

In E2, cued motor adaptation was similar as in E1. The main differences were the number of learning blocks and the timing of the trials. This time only one learning block (162 trials) was used instead of two learning blocks (2 times 81 trials). One trial of cued adaptation experiment (E2) took exactly 4.5 seconds. First, participants performed a target reach. Immediately after the reaching, extra fixation time was added in order to obtain exactly 4.5 seconds per trial. After 4.5 seconds, the next trial started automatically. If participants exceeded 3 seconds for the reaching movement, a warning sign was shown and a high pitch was played in order to instruct participants to speed up. The correct reaching time was between 175 ms and 375 ms, which was indicated with a target color change. If reaching time was above 375 ms, the target became purple. If reaching time was below 175 ms, the target became red. Two breaks of 1 min were given to participants, one before the 4^th^ cycle (trial 36) of the baseline and one before the 9^th^ cycle (trial 81) of the learning block.

#### Preregistration

The flanker dual-task (E1) study was preregistered online: http://aspredicted.org/blind.php?x=wj3im4. This preregistration included the main hypotheses, the key dependent variables, the amount of participants, the main analyses and some of the secondary analyses investigated. In the preregistration, we mentioned that we would include 30 participants per group. However, in the end, we included 21 additional subjects (11 young, 10 older adults) for this preregistered study (E1) in the context of a master student project but these additional participants did not change the outcome of our study.

The main pre-registered analysis tested for a significant negative correlation between explicit adaptation and dual-task cost.

#### Cognitive assessment

##### Visuospatial working memory capacity

During the flanker dual-task experiment (E1), participants performed a working memory task. The task is the same as described in Christou et al., 2016, but is slightly different from the working memory dual-task experiment (E2), described above. The main differences with the task of E2 were: 1) Only three, four, five or six red circles (0.8 cm diameter) were used (the two target condition was not used); 2) The timing of the events was the same in both task, except that, here, participants had three seconds to indicate the right answer instead of two seconds; 3) Indicating the right answer was performed by making a button press, instead of moving the cursor to a “yes answer” location or a “no answer” location; 4) After the three seconds for responding, they had to fixate on a small blue cross (0.2 cm x 0.2 cm) for one second; 5) In total, one trial had a fixed time duration of nine seconds, instead of eight seconds. All participants had a first practice session of eight trials. After this, each participant had to complete 40 trials instead of 50 trials as in experiment E2. The 40 trials contained three, four, five or six red circles (10 trials/condition) with all conditions randomly mixed.

## Data availability

All data, analysis scripts and supplementary materials are available on Open Science Framework: https://osf.io/ks2j8/

## Data analysis

Analyses of E1 were preregistered, while analyses of E2 were performed without preregistration.

### Preprocessing

All analyses and statistical calculations were performed in MATLAB 2018b (The MathWorks). For each reaching movement, the hand angle (relative to target angle) was calculated from the first data point exceeding 4 cm distance from the middle of the starting point. The time for reaching 4 cm was on average 172 ms in E1 and 144 ms in E2. The hand angle was the primary dependent variable in cued adaptation experiments. The angular error is the angle the cursor deviated from the target. Angular errors above 60 ° were due to inattentive reaches to previous target directions and were considered as outliers. These outliers were removed before processing the data. The number of outliers that we removed, constituted 0.81 % of all trials for E2. For E1 this procedure was not implemented but this would not change the outcome of the study. The statistically significant threshold was set at p<0.05 for the ANOVA’s. We reported effect sizes (eta squared: *η*^2^) as well as F and p-values.

### Analysis 1: Final adaptation level

To assess the final adaptation level of each participant, we averaged the hand angles of the last 18 trials of the learning block. These hand angles were first corrected with the average hand angles of the last 18 baseline trials before each learning block. Statistical comparison was performed with a 2-way ANOVA with two between-subject factors: age group and rotation direction.

### Analysis 2: Implicit adaptation

In E1, we applied the same analysis as described in Vandevoorde and Orban de Xivry (2019). The first adaptation block was corrected for baseline errors by subtracting the average error of the last 18 trials of baseline. The second adaptation block was corrected by subtracting the average error of the last 18 trials of washout. We analyzed the data in all the uncued trials that were preceded by a cued trial (nine uncued trials per learning block). The amount of implicit learning was calculated per learning block as the average of the uncued trials (Morehead et al., 2015). For each learning block separately, we performed a 2-way ANOVA with the implicit adaptation level as dependent variable and with age and rotation direction as between-subject factors.

In E2, the adaptation block of cued adaptation was corrected for baseline errors by subtracting the average error of the last 18 trials of baseline. We analyzed the data in all the uncued trials of the learning block (18 uncued trials). The amount of implicit adaptation was calculated as the average of all uncued trials. One 2-way ANOVA was used, with the between-subject factors, age and rotation direction, and with the implicit adaptation as dependent variable.

### Analysis 3: Explicit adaptation

In E1, we applied the same analysis as described in Vandevoorde and Orban de Xivry (2019b). The amount of explicit learning was calculated by subtracting hand direction in the uncued trials (Analysis 2) from the cued trials immediately preceding those (Morehead et al., 2015). Two separate 2-way ANOVA’s were used to analyze the first and second learning block with the explicit adaptation level as the dependent variable and with the between-subject factors, age and rotation. The 2-way ANOVA to analyze the first learning block of experiment E1 was preregistered as a primary analysis.

In E2, baseline subtraction of learning block was the same as in Analysis 2. The amount of explicit learning was calculated by subtracting hand direction in the uncued trials from the cued trials immediately preceding those (Morehead et al., 2015). A 2-way ANOVA was used to analyze the learning block for cued adaptation with the explicit adaptation level as the dependent variable and with the between-subject factors, age and rotation.

### Analysis 4: Dual-task cost measures

In the flanker dual-task, the dual-task cost (DTC) is calculated for each subject in two different ways based on median reaction time (RT) of reaching (motor-task) and the median reaction time of the flanker task (cognitive-task). The first dual-task cost was preregistered:

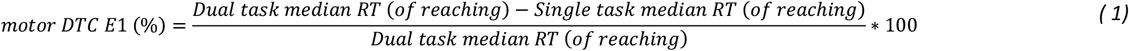

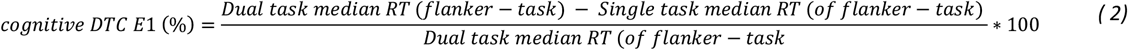

In the working memory dual-task design, the dual-task cost is calculated for each subject in two different ways based on reaction time of reaching (motor-task) and WMC (cognitive-task):

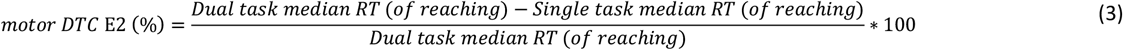

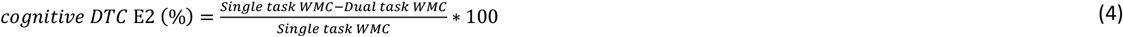

We used two different formulas for calculating dual-task costs, derived from single-task (ST) and dual-task (DT) performance: DTC = [(DT-ST)/DT]*100 if an increase in the metric was related to a performance decline (e.g. higher reaction time) and DTC = [(ST-DT)/ST]*100 if an increase in the metric was related to a performance increase (e.g. higher working memory capacity). Positive and negative DTC values therefore always corresponded with, respectively, decreased and increased performance from single-to dual-task condition (Doumas et al., 2008; Boisgontier et al., 2013).

Two-sample two-tailed t-tests were applied to verify whether these dual-task costs were different for young and old participants. We also compared dual-task costs with zero reference via one-sample t-tests against zero. We reported effect sizes (cohen’s d) as well as t and p-values for all t-tests.

### Analysis 5: Correlation dual-task cost and explicit adaptation

Robust linear regression (robustfit in Matlab) was executed between the measure of explicit adaptation and dual-task cost. Robust linear regression was performed in order to verify that correlations were not influenced by between group differences in the variables. Explicit adaptation (Y) was estimated using a linear combination of dual-task cost (X), a binary age vector (G) and the interaction of X and G in the regression equation with intercept A and regression coefficients (B, C, D):

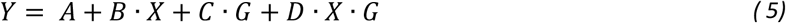

Standardized beta coefficients (B, C, D) by first converting variables X and Y to z-scores.

Our main prediction was that, especially in elderly people, the two variables (explicit adaptation and dual-task cost) will be negatively correlated. This robust linear regression was preregistered for the flanker dual-task experiment with the motor dual-task cost E1 (Eq. 1).

### Analysis 6: Working memory capacity (WMC)

The computer-based working memory task allows the determination of WMC with the K-value, estimating the number of items that can be stored in working memory (WM) (Vogel et al., 2005), calculated as K = S(H-F) using two (in E2) or three (in E1) to six items. This is similar to the original experiment (Vogel et al., 2005) but differs from what previous adaptation studies (Christou et al., 2016) have used where the K-value (i.e. K56) was obtained from the trials with five and six items only. We chose to measure WMC with all items because it allowed us to better estimate the K-value for elderly than with the K-value with only five and six items. This correlation with working memory capacity measured with all items and overall adaptation was also significant in the study of Christou et al. (2016) (personal communication from Dr. Galea). Two-sample two-tailed t-tests were applied to verify whether the working memory capacity was different for young and old (preregistered as a secondary analysis for E1). Working memory capacity was related to the level of explicit adaptation. Robust linear regression (robustfit in Matlab) was performed in order to partial out the effect of age group on the correlations as explained in Analysis 5.

## Results

It is well established that explicit adaptation is reduced in older adults (Heuer and Hegele, 2008; Vandevoorde and Orban de Xivry, 2019). One hypothesis to explain this reduction is the *cognitive resources hypothesis* that states that older adults require more cognitive resources compared to younger adults for unperturbed reaches which leaves then fewer cognitive resources available for the adaptation process. To assess the amount of cognitive resources used for reaching movements, we introduced a cognitive task during the baseline period of a visuomotor rotation experiment, which allowed us to estimate the amount of cognitive resources used to reach to targets via the dual-task costs. The higher the dual-task costs the more cognitive resources were required for unperturbed reaching.

### Flanker dual-task design (E1): Decrease in the explicit component of adaptation with age

To quantify the overall effect of age on motor adaptation, we measured its extent by looking at hand angles over the last 18 trials of the first learning block (Analysis 1). We observed a decrease of final adaptation level for older adults in the cued visuomotor rotation experiment (Figure 3A-B; learning: F(77) = 8.5, p = 0.005, ɳ^2^= 0.1; relearning: F(77) = 11.1, p = 0.001, ɳ^2^= 0.1). We observed no difference between younger adults and older adults for implicit adaptation (Figure 3C: Analysis 2; Learning: F(1,77) = 3.1, p = 0.08, ɳ^2^=0.03; Relearning: F(1,77) = 0.3, p = 0.60, ɳ^2^=0.003). One outlier data point was present in the young adult’s group in the first learning block for implicit adaptation (Figure 3C). However, removing this outlier would not change the result for implicit adaptation during the first learning block. Explicit adaptation was decreased for older adults in both learning blocks (Figure 3D; Analysis 3; Learning: F(1,77) = 4.4, p = 0.04, ɳ^2^=0.05; Relearning: F(1,77) = 9.5, p = 0.003, ɳ^2^= 0.10).

**Figure 3:**
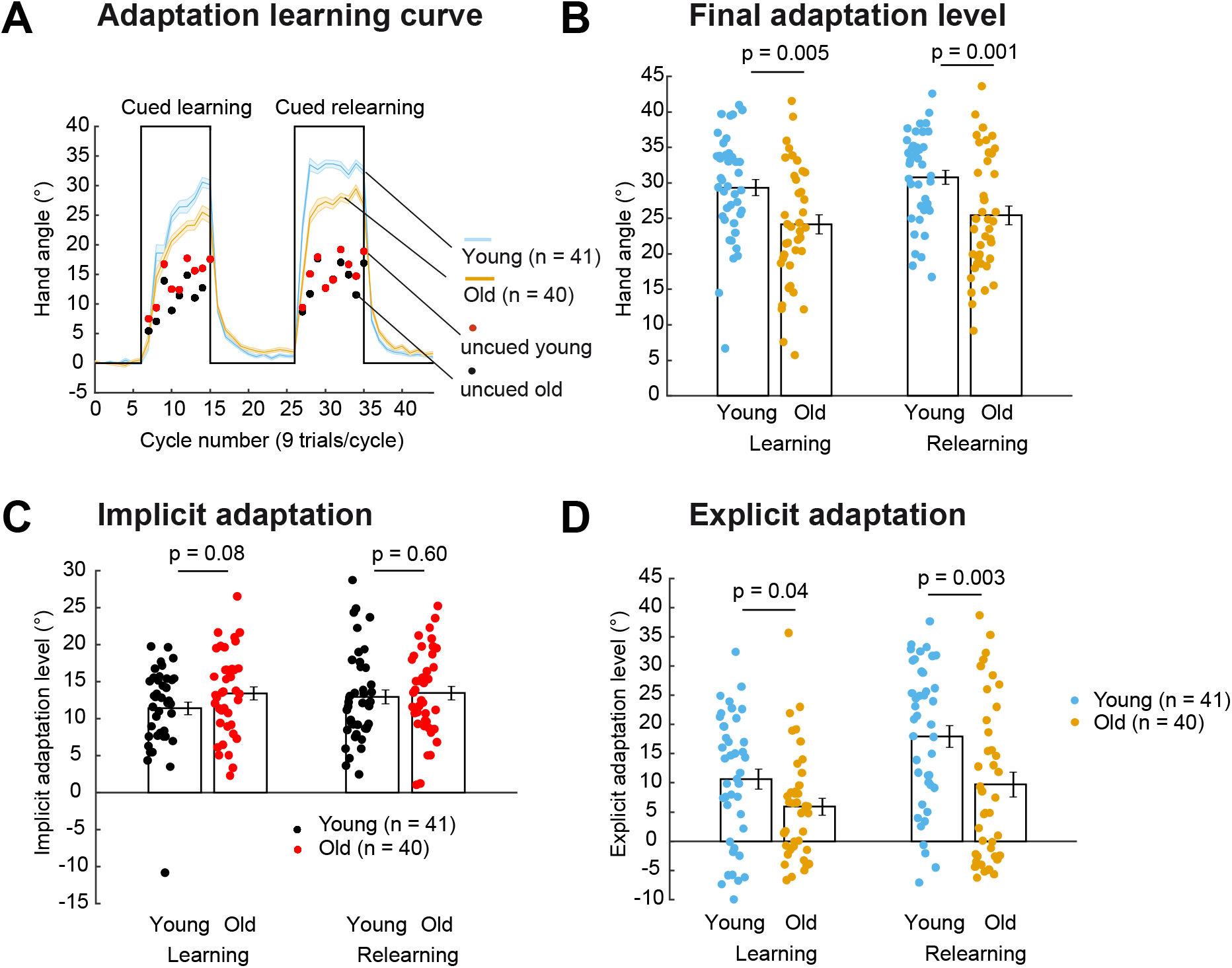
Differences in motor adaptation between young and older adults. **A)** Decreased overall cue-evoked adaptation in older adults compared to younger adults in learning and relearning block. During the uncued trials, the level of implicit adaptation was measured and the cued trials preceding the uncued trials allowed us to calculate explicit adaptation. **B)** Final adaptation level at the end of the learning and relearning block was lower in older adults. **C)** Implicit adaptation was not different for younger adults and older adults in learning and relearning. **D)** Explicit adaptation was reduced in the learning and the relearning block for older adults compared to younger adults.

### Flanker dual-task design (E1): No evidence for the cognitive resources hypothesis

In a first preregistered study, the flanker task was introduced as the cognitive task during the baseline period in order to assess dual-task costs. According to the *cognitive resources hypothesis*, we expected a larger dual-task cost in elderly people and a negative correlation between explicit adaptation and the dual-task costs.

The motor dual-task cost (Eq. 1) was not different between young and older adults (Figure 4A, Analysis 4; t(79) = −1.6; p = 0.12, d = 0.35), which could indicate that the amount of cognitive resources required for unperturbed reaching was not higher for older adults. In addition, we observed no difference between young and older adults for the cognitive dual-task cost (Eq. 2) (Figure 4B: Analysis 4; t(79) = 0.4; p = 0.66, d = 0.10). These two dual-task costs measures (Eq. 1 & 2) were positive, for both age groups (Figure 4A-B; Motor DTC(1): Young: t(40) = 6.1, p = 4×10-7, d = 0.95, Old: t(39) = 7.5, p = 5×10-9, d = 1.18; Cognitive DTC: Young: t(40) = 7.7, p = 2×10-9, d = 1.21, Old: t(39) = 4.9, p = 2×10-5, d = 0.78). These positive dual-task costs show that the dual-task manipulation worked as the performance of the motor and cognitive tasks was reduced in dual-task condition compared to single-task condition.

**Figure 4:**
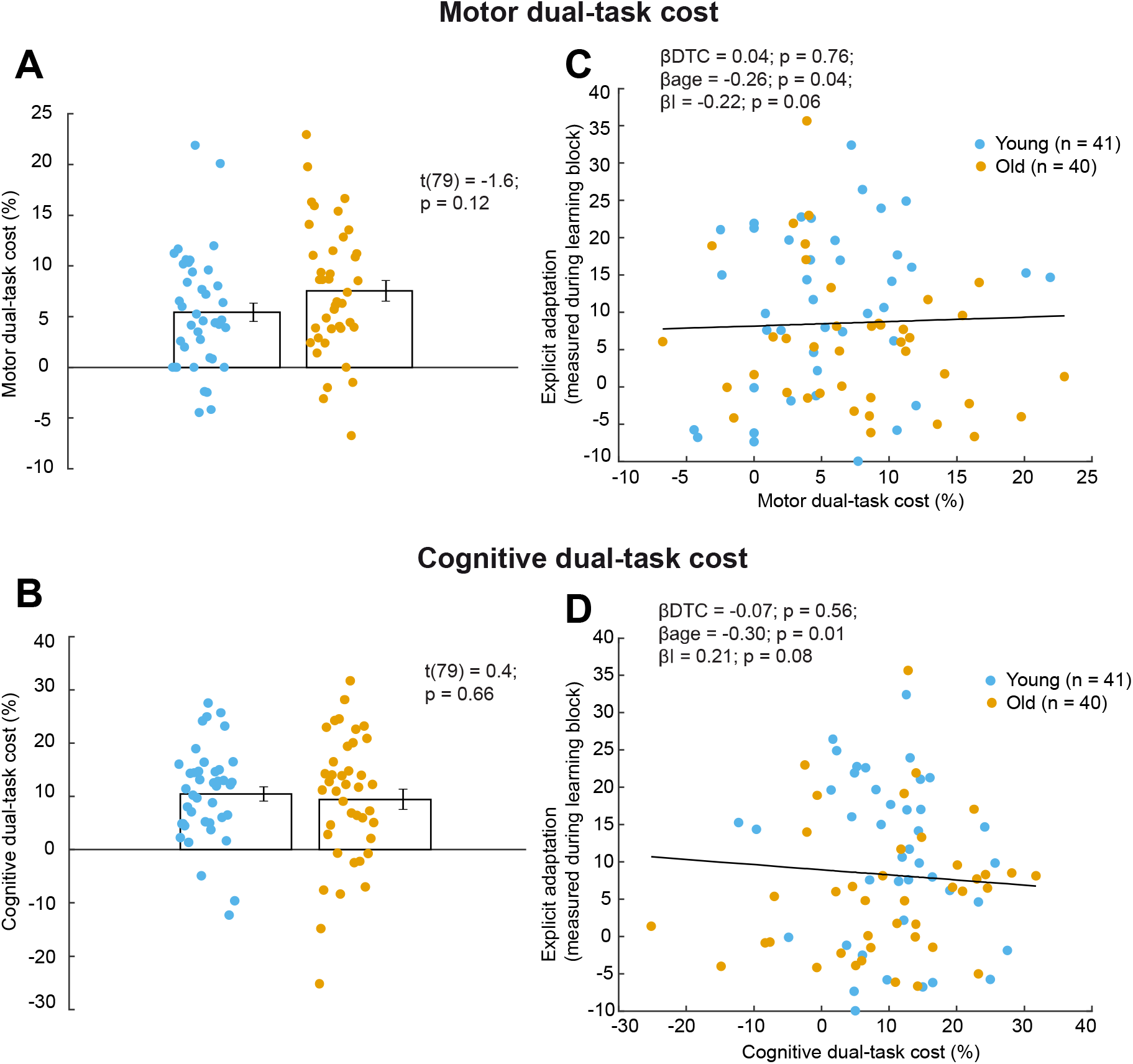
Motor and cognitive dual-task costs: **A-B)** Dual-task costs were not different for young and older adults. However, the two dual-task costs were bigger than zero for both age groups. **C-D)** No link was observed between explicit adaptation and dual-task costs. DTC, dual-task cost; I, interaction.

In addition, we did not find any evidence in favor of the cognitive resource hypothesis as we observed no link between the preregistered motor dual-task cost and explicit adaptation (Figure 4C, Analysis 5; β = 0.04; p = 0.76). In addition, no relation was observed between the cognitive dual-task cost and the amount of explicit adaptation (Figure 4D; Analysis 5; β = −0.07; p = 0.56). As such, we failed to find any evidence for the cognitive resources hypothesis.

### Flanker dual-task design (E1): Link between working memory capacity and explicit adaptation

However, we also quantified working memory capacity (Analysis 6), because a high working memory capacity was linked to an individual’s capacity to use an explicit strategy (Christou et al., 2016). Working memory capacity was lower for older adults (Figure 5A: t(76)= 4.46; p = 2.8×10-5, d = 1.01). Furthermore, there was a positive link between explicit adaptation and working memory capacity (Figure 5B: β_WMC_ = 0.39; p = 0.002). This link was also observed for the amount of explicit adaptation in the relearning block (β_WMC_ = 0.39; p = 0.004).

**Figure 5:**
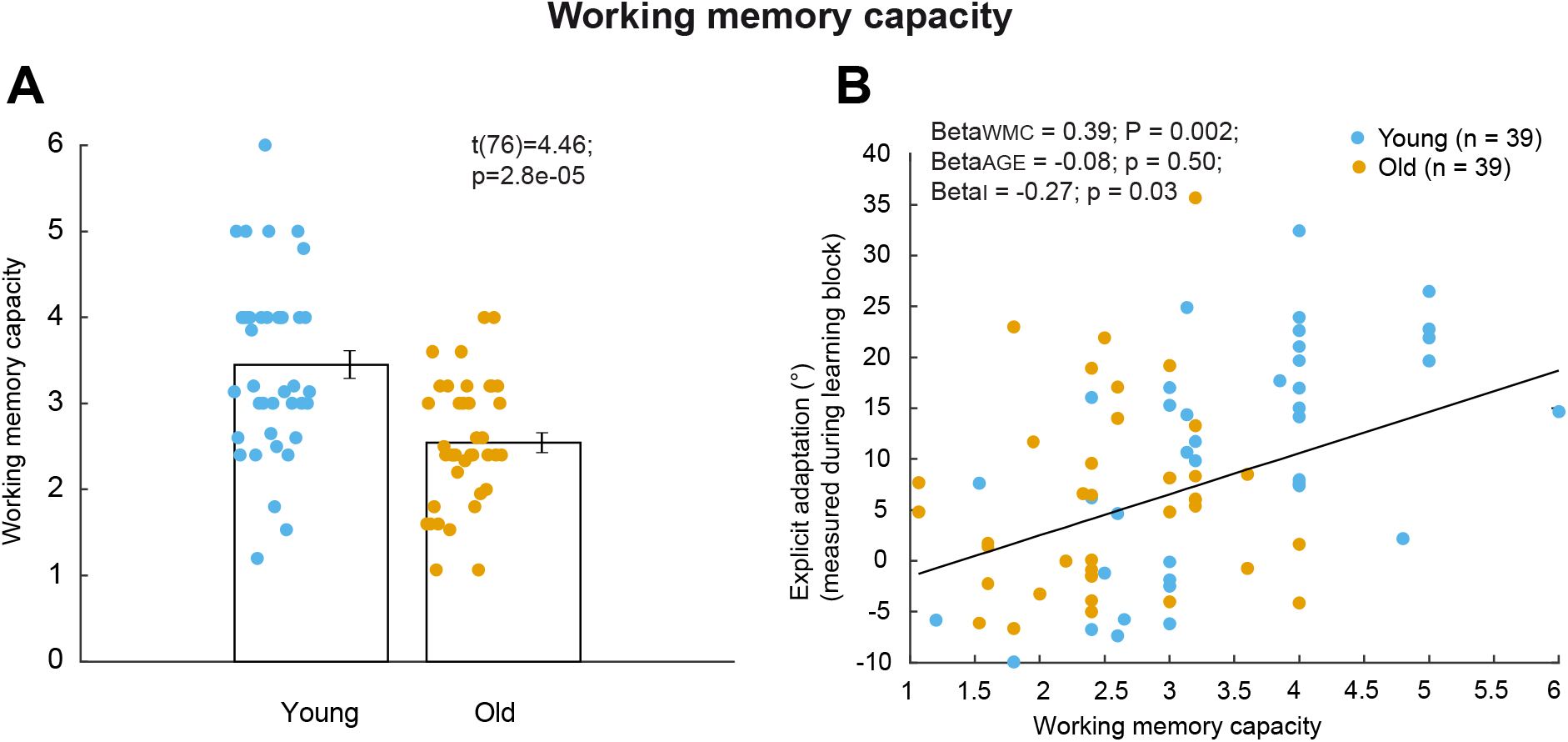
Link between working memory capacity (WMC) and explicit adaptation. **A)** Working memory capacity was lower for older adults compared to younger adults. For three (of 81) subjects no working memory capacity data was obtained. **B)** A positive correlation existed between explicit adaptation during the learning block and working memory capacity.

One disadvantage of the flanker dual-task was the serial design of the reaching and cognitive tasks. Since the same cognitive resources could be used for the sequential tasks, the dual-task costs might not be the best estimate of the amount of cognitive resources used for simple reaching movements (despite the fact that there was a dual-task cost). Therefore, we used a second cognitive-motor dual-task design where a working memory task was performed in parallel to the reaching task. The working memory task appeared suitable as we just showed that it is linked to the explicit component of adaptation.

### Working memory dual-task design (E2): Decrease in the explicit component of adaptation with age

In terms of adaptation, the results were largely similar to the results obtained in the first experiment. We observed a decrease of final adaptation level for older compared to younger adults in the cued visuomotor rotation experiment with a single learning block (Figure 6A-B; Analysis 1; F(1,58) = 9.7, p = 0.003, ɳ^2^ = 0.14). Implicit adaptation was slightly larger in old compared to young participants (Figure 6C; Analysis 2; implicit adaptation: F(1,58) = 1.6, p = 0.22, ɳ^2^ = 0.02), while explicit adaptation was decreased for older adults (Figure 6D; Analysis 3; explicit adaptation: F(1,58) = 10.3, p = 0.002, ɳ^2^ = 0.14).

**Figure 6:**
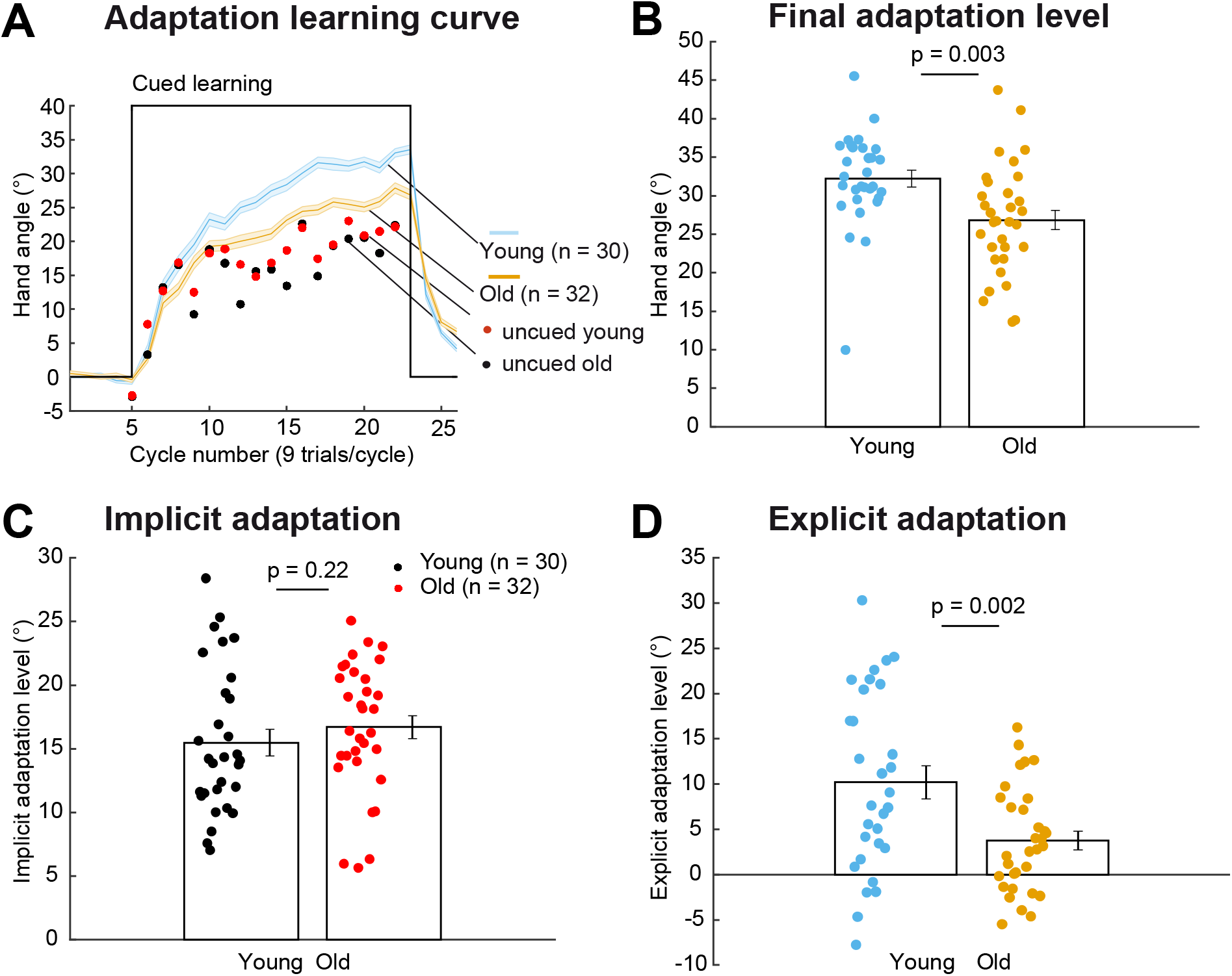
Differences in adaptation between young and older adults. **A)** Decreased overall cue-evoked adaptation in older adults. During the uncued trials, the level of implicit adaptation was measured and the cued trials preceding the uncued trials allowed to calculate explicit adaptation. **B)** Final adaptation level at the end of the learning block was lower in older adults. **C)** Implicit adaptation was not different for younger and older adults. **D)** Explicit adaptation was reduced for older adults compared to younger adults.

### Working memory dual-task design (E2): No evidence for the cognitive resources hypothesis

In the working memory dual-task design, the working memory capacity task was implemented as a cognitive task during the baseline period (Figure 2B) in order to assess dual-task costs during unperturbed reaching movements. The hypothesis remained unchanged from the first preregistered study: we expected a negative link between two dual-task costs and explicit adaptation according to the *cognitive resources hypothesis*.

The dual-task cost variables were calculated as the relative difference in median reaction time for reaching (Eq. 3) and as the relative change in working memory capacity (Eq. 4) from single to dual-task condition (Analysis 4). The motor dual-task cost (Eq. 3) was not different between young and older adults (Figure 7A: Analysis 4: t(60) = 1.4, p = 0.18, d = 0.35). In contrast to our hypothesis, the cognitive dual-task cost (Eq. 4) was not different between younger and older adults (Figure 7B: Analysis 4: t(60) = −0.78, p = 0.44, d = 0.20). The motor dual-task cost (Eq. 3) was not different from zero (Figure 7A: Young: t(29) = 1.2, p = 0.25, d = 0.21; Old: t(31) = −0.65, p = 0.52, d = −0.11). However, the cognitive dual-task cost (Eq. 4) was positive, both for young and old (Figure 7B: Young: t(29) = 4.2, p = 0.0002, d = 0.76; Old: t(31) = 4.1; p = 0.0003, d = 0.72). This positive dual-task cost indicates that unperturbed reaching partially depends on working memory capacity resources for both young and older adults. In addition, it showed that the dual-task manipulation worked. However, contrary to our hypothesis, older adults did not require more of these resources than younger adults did.

**Figure 7:**
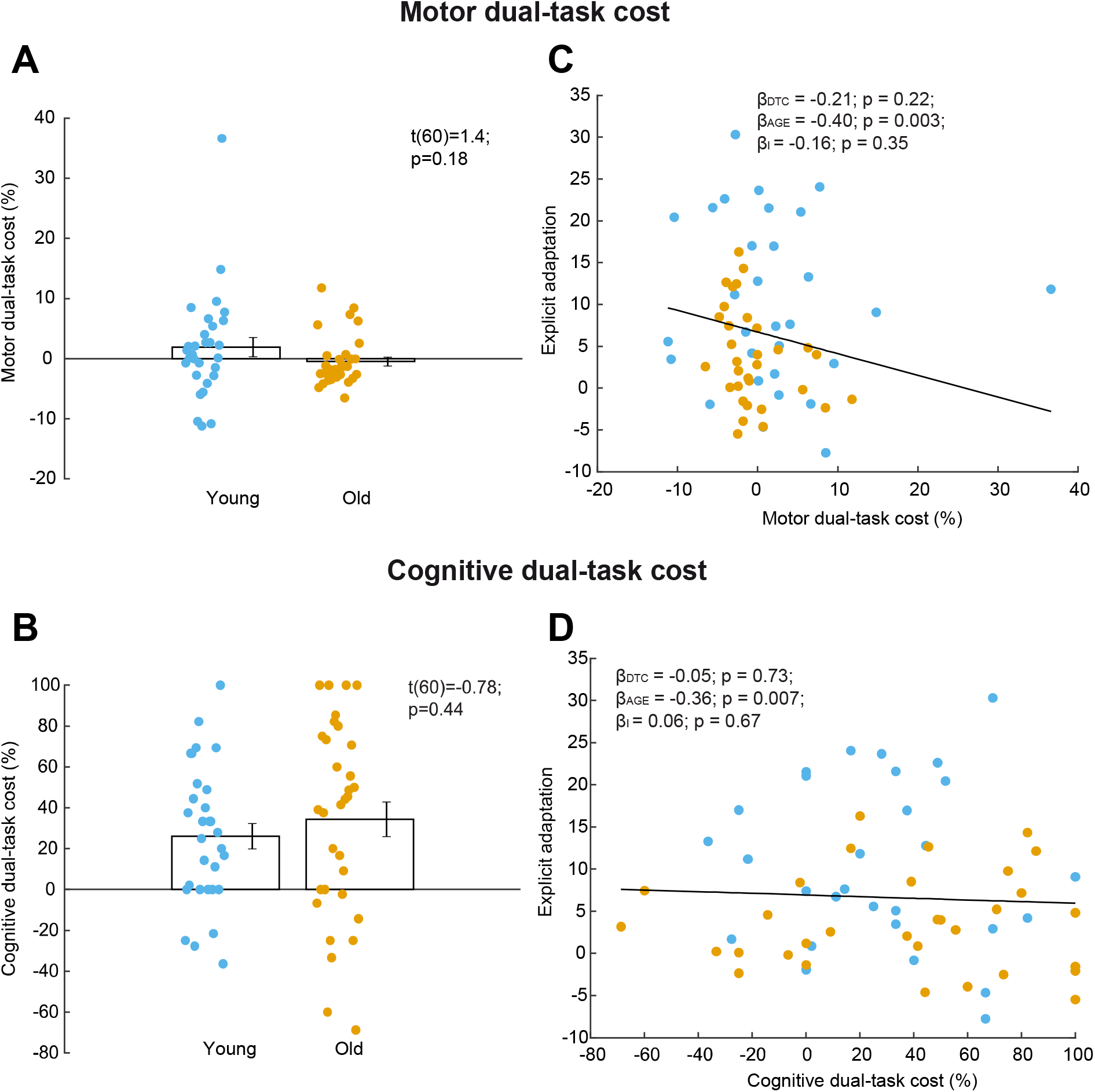
Motor and cognitive dual-task cost. **A)** Motor dual-task cost calculated as the relative change in median reaction time of reaching was not different between young and older adults. **B)** Cognitive dual-task cost for working memory capacity was not different between young and older adults. **C)** No link was observed between explicit adaptation and motor dual-task cost. **D)** No link between explicit adaptation and cognitive dual-task cost.

Finally, there was neither a link between motor dual-task cost (Eq. 3) and explicit adaptation (Figure 7C: Analysis 5; β = −0.21; p = 0.22) nor a link between cognitive dual-task cost (Eq. 4) and explicit adaptation (Figure 7D: Analysis 5; β = −0.05; p = 0.73). One outlier appears to be present in Figure 7B-C, however, given the use of robust regression methods, the outlier does not cause the absence of correlation. Therefore, we did not find any evidence for the cognitive resources hypothesis. The absence of any robust link between other dual-task cost measures and explicit adaptation in the flanker dual-task (Figure 4; Supplementary Table 1) and in the working memory dual-task (Figure 7; Supplementary Table 2) shows that it is unlikely that a link exists between the cognitive resources required for unperturbed reaching in baseline and the level of explicit adaptation.

### Working memory dual-task design (E2): Link between working memory capacity and explicit adaptation confirmed

In this experiment, working memory capacity, which was quantified in the single- and dual-task condition, was lower in older adults (Figure 8A-B: Analysis 6: t(60) = 2.6, p = 0.01, d = 0.67; t(60)= 2.6, p = 0.01, d = 0.67). As a confirmation of our earlier finding in experiment E1, we found a link between explicit adaptation and working memory capacity in the single-task condition (Figure 8C: Analysis 6: β = 0.29; p = 0.03). This link did not reach significance in the dual-task condition (Figure 8D: Analysis 6: β = 0.18; p = 0.18).

**Figure 8:**
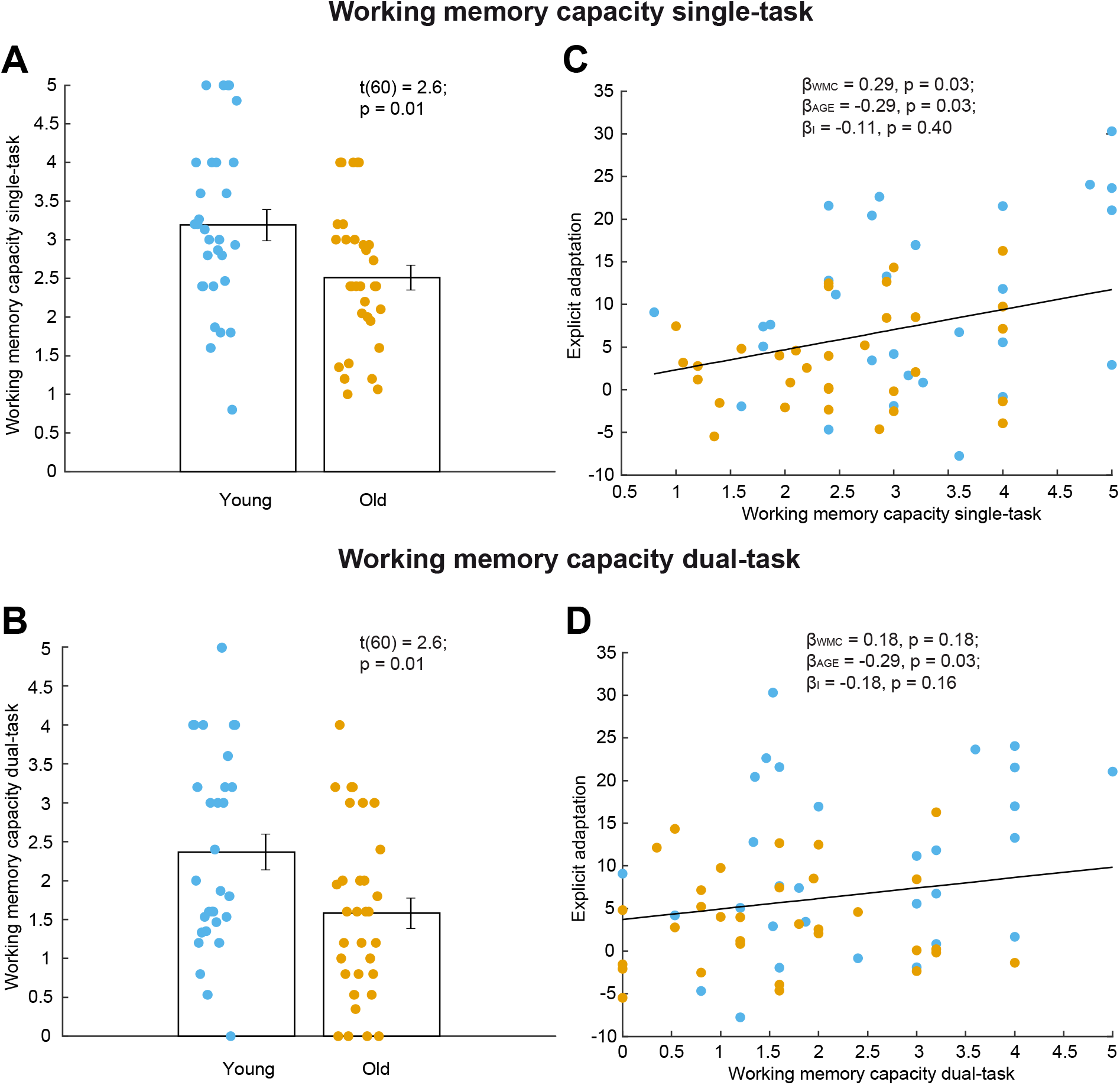
Link between working memory capacity and explicit adaptation. **A)** Single-task working memory capacity was lower for older adults. **B)** Dual-task working memory capacity was lower for older adults. **C)** Positive link between explicit adaptation and single-task working memory capacity. **D)** No link observed between explicit adaptation and dual-task working memory capacity.

To conclude, we found no evidence for the cognitive resources hypothesis as none of the cognitive or motor dual-task cost measurements in baseline were related to explicit adaptation for both dual-task designs. In addition, cognitive and motor dual-task costs were not different for young and older adults. However, in both experiments, working memory capacity was consistently linked with explicit adaptation and working memory capacity was smaller in older adults.

### Link between cognitive status and explicit adaptation: exploratory analyses

Finally, several other variables were affected by the dual-task condition as indicated by positive cognitive and motor dual-task costs (Supplementary Table 1 & Table 2). In addition, a negative link was observed between reaction time flanker task and explicit adaptation (Supplementary Table 1, single-task condition: β = - 0.50; P = 0.01; dual-task condition: β = - 0.49; P = 0.003) and between explicit adaptation and reaching speed errors (Supplementary Table 1, β = −0.43; P = 0.003). This might indicate that participants with increased choice reaction time have more difficulties to develop an explicit strategy. In a large-scale longitudinal study with 1265 volunteers ranging from age 17 to 96, reaction time increased with age, which might reflect a general age-related slowing (Fozard et al., 1994). Also the reaching speed errors were mainly caused by too low motor speed. Therefore, age-related movement slowing (Birren and Fisher, 1995; Salthouse, 2000; Seidler et al., 2010) might be limiting explicit strategy as well. On the other hand, increased choice reaction time in the flanker task might be resulting from a decreased selective attention (Kopp et al., 1994). Finally, a positive link with the figure recall score (Supplementary Table 4, β = 0.44; P = 0.0004) can be interpreted as visuospatial abilities or visuospatial memory being important for explicit strategy. The links between these other cognitive variables (reaction time flanker task, reach speed errors and figure recall) and explicit adaptation have never been reported but are interesting to test more extensively.

## Discussion

In the present study, we did not find any evidence that the amount of explicit adaptation was linked to the total amount of cognitive resources minus those used during unperturbed reaching. In both cognitive-motor dual-task experiments, cognitive and motor dual-task costs measured during the baseline period were similar for young and older adults, indicating that the amount of cognitive resources used during unperturbed reaching is not different for these groups. Altogether, we found no evidence for the cognitive resources hypothesis (Figure 1). However, associations between the level of explicit adaptation and between visuospatial working memory, independent of age, in two experiments together pointed towards the importance of cognitive abilities for the achieved level of explicit adaptation. In conclusion, it was rather the reduced working memory abilities of older adults and not the saturation of cognitive resources in unperturbed reaching movement that is linked to the reduced explicit adaptation in older adults.

However, this does not resolve the contradiction that existed between previous motor tasks where elderly exhibit increased cognitive control to improve motor performance (Heuninckx et al., 2008) and the age-related decline of the cognitive component in motor adaptation. One explanation is that increased cognitive control was present only at lower cognitive load, while at higher load it failed to be useful. Indeed, increased prefrontal cortex activation in older compared to younger adults was limited to low levels of working memory load. While at higher loads, working memory performance was lower in older adults and the increased brain activity disappeared (Mattay et al., 2006; Schneider-Garces et al., 2009; Cappell et al., 2010).

The idea of reduced working memory contributing to decline of motor adaptation is in line with Anguera et al. (2011). They suggested that spatial working memory contributes to the age-related deficits in visuomotor adaptation. They observed a link between a measure of visuospatial working memory and early rate of learning. However, they note that the observed link would fail after correction for multiple comparisons. We expanded their work by dissociating overall adaptation in an implicit and an explicit component of adaptation, of which only the explicit component is known to be affected with aging (Vandevoorde and Orban de Xivry, 2019). In addition, we quantified visuospatial working memory with a paradigm that allowed establishing the link with the explicit component (Christou et al., 2016). By these adjustments together with bigger sample sizes, a robust link could be observed between working memory capacity and the explicit component in two experiments, which remained significant after correcting for age (Figure 5B, Figure 8C). In addition, older adults’ working memory capacity was lower than younger adults’ capacity in both experiments (Figure 5A, Figure 8A-B). In addition, recently Rajeshkumar and Trewartha (2019) showed that, when reducing working memory demands during motor adaptation by repeating a specific order of target locations, the difference between young and older adults’ adaptation rates disappeared. Together, this allows us to confirm that working memory abilities seem to have an important contribution to the age-related deficits of motor adaptation.

Since working memory appears to be robustly linked with explicit strategy, cognitive training might resolve the age-related decline of motor adaptation. It would be interesting to verify whether specifically training visuospatial working memory has an impact on the level of explicit strategy. This approach has been attempted by Anguera et al. (2012) in younger adults, however, without success. They even mention an opposite effect after extensive training of visuospatial working memory: The training might have resulted in depletion of spatial working memory resources, which negatively affected subsequent visuomotor adaptation. However, it appears that availability of working memory resources can modulate the rate of adaptation in younger adults. Nevertheless, for older adults the outcome might still be beneficial since working memory training is effective in elderly and might be a useful tool for cognitive intervention (Klingberg, 2010; Karbach and Verhaeghen, 2014). However, such training study should be carefully designed given the limitations in working memory training literature such as lack of consistency in experimental methods and findings (Morrison and Chein, 2011). Since correlation does not imply causation, it is recommended to design studies that verify causality between working memory and explicit adaptation.

Our understanding of the cognitive component of motor adaptation is only at an early phase. Recently, a link between working memory and explicit strategy has been revealed (Christou et al., 2016). Afterwards, this link with working memory has been further explored in McDougle and Taylor (2019). They showed that two different representations of working memory are involved in generation of explicit strategy: one is parametric mental rotation and the other is discrete response caching. With lower number of targets (2), response caching is the main working memory process involved. With a higher number of targets (12), both working memory processes seem to be involved: at early learning mental rotation is mainly involved, later in learning a switch to response caching is possible (McDougle and Taylor, 2019). Only three targets were used in our learning blocks, which might suggest that response caching could be involved in our measure of explicit strategy. Our working memory capacity measurement was similar to response caching in the sense that participants had to remember a discrete number of items (Spellman et al., 2015). The link that we observed between working memory capacity and explicit strategy is therefore originating from the involvement of response caching. The link that Anguera et al. (2011) observed between working memory and visuomotor learning originates from mental rotation which involves the mental rotation of objects to indicate which ones are similar. Therefore, together we show that both aspects of working memory, response caching and mental rotation, might be contributing to the age-related declines of motor adaptation. It would be interesting to further explore this idea with the recent approaches taken in McDougle and Taylor (2019) by constraining reaction time and applying small vs large set sizes. However, careful design is important since reaction times tend to be much higher in older adults and are not a good indicator for explicit strategy in older adults (Vandevoorde and Orban de Xivry, 2019).

Working memory capacity is often associated with dorso-lateral prefrontal cortex changes (Curtis and Esposito, 2003). Age-related changes in dorsolateral prefrontal cortex activity were observed during information retrieval from working memory (Rypma and D’Esposito, 2000). Working memory impairment in normal aging might originate from a deficit in top-down suppression deficit of task-irrelevant representations (Gazzaley et al., 2005). Older adults have deteriorated frontal lobe function and structure (Moscovitch and Winocur, 1995; Raz and Rodrigue, 2006). These age-related brain changes might be involved in the age-related decline of explicit adaptation. In addition, working memory declines with aging (Jost et al., 2011) which might also contribute to the decline of explicit adaptation (Anguera et al., 2011). Therefore, we predict to see different neural activation in prefrontal cortex depending on the level of explicit strategy acquired. A relationship between neural activity in dorsolateral prefrontal cortex and early learning rate was indeed observed in Anguera et al. (2011). However, replication of this relation is useful to verify robustness. In addition, it would be interesting to further refine the region of interest, quantify explicit strategy directly, and disturb activity of this brain region to investigate causality.

The goal of both cognitive-motor dual-task designs was to quantify the amount of cognitive resources applied during unperturbed reaching movement. This is a common procedure in gait (Ebersbach et al., 2011) and balance studies (Melzer et al., 2001; Huxhold et al., 2006); however, for reaching movements it has been reported for only a few studies (Bekkering et al., 1994; Pratt and Neggers, 2008; Ma et al., 2009). The presence of motor and cognitive dual-task costs shows that performance of both tasks was negatively affected by performing them simultaneously. This negative impact might indicate that the same cognitive resources were required for both tasks and that the dual-task design succeeded in its goal of measuring cognitive resources during reaching. For instance, a reduction of working memory capacity with more than 20 % was observed when combining reaching with the working memory capacity task; working memory capacity resources appear therefore to be involved in unperturbed reaching. Dual-task costs are assumed to be an estimation for the amount of cognitive resources applied (Boisgontier et al., 2013). In our experiments, we did not observe differences in dual-task cost between young and older adults. However, it is likely that increasing the complexity of the dual-task by reducing the time per trial or by asking to remember a higher number of red dots during the working memory task (Figure 2) would result in different dual-task cost for young and old participants. This would be in line with studies that reported a more pronounced effect of aging, when the cognitive load of a cognitive-motor dual-task was higher (Boisgontier et al., 2013). Nevertheless, a relation between dual-task cost and explicit adaptation seems unlikely given the absence of any relation so far (Figure 4C-D, Figure 7C-D).

In both the cognitive-motor dual-tasks and the adaptation experiments, a trial-by-trial reward was provided to participants upon successful task performance. This reward was implemented to motivate participants throughout the duration of the experiment. However, this reward-feedback makes it impossible to dissociate performance driven by errors from performance driven by reward (Cashaback et al., 2017). Huang et al. (2018) recently showed that reward modulated adaptation learning rates positively and similarly in both young and older adults. Therefore, reward might not have a differential effect in our results. However, additional experiments are required with and without reward-feedback to confirm this. Reward might also affect the relation between working memory capacity and explicit adaptation. The working memory assessment in the flanker dual-task experiment was without reward-feedback, while during working memory dual-task a reward was implemented for working memory trials.

In the flanker dual-task, dual-task costs were observed for the motor and the cognitive task (Figure 4A-B). Therefore, priority is not given to one of the tasks. In the working memory dual-task, dual-task costs were only observed for the cognitive task (Figure 7A-B), which might indicate that priority is given to the motor task. However, if inspecting other dual-task costs (Supplementary Table 1), it becomes apparent that reaching was slower and more reaching errors occurred during the dual-task condition. Therefore, the reaching was affected as well in the working memory dual-task and no priority was given to one of the tasks.

The timing was different for the two dual-task experiments. A limitation of the flanker dual-task was the serial design of the cognitive and motor task. Therefore, it resembled more a switching-task instead of a dual-task (Monsell, 2003). To tackle this limitation, we introduced the second dual-task with true parallel features. However, even for the parallel design, no relation was observed between explicit adaptation and dual-task costs. The cognitive tasks introduced in the baseline were different, but single-task performance for both cognitive tasks appeared to be linked to explicit adaptation.

The first motor adaptation paradigm consisted of two learning blocks while the second paradigm contained only one learning block. In addition, individual trials were strictly constrained in timing in the second adaptation paradigm but not in the first paradigm. This might have resulted in different explicit adaptation levels. However, a difference of explicit adaptation levels for both age groups was present in both experiments, although maybe more pronounced in the second paradigm as suggested by the higher effect size (E1: ɳ^2^=0.05, E2: ɳ^2^ = 0.14).

Besides shared cognitive resources among the cognitive and motor tasks, an increased attentional cost might explain why performance is reduced in dual compared to single-task (Wickens, 1980). Therefore, the cognitive dual-task cost in our working memory dual-task might reflect a divided attention between the motor and cognitive tasks instead of an involvement of working memory resources in the unperturbed reaching behavior. In addition, studies often link dual-task performance to other components of the executive system, such as planning, shifting, inhibition or coordination (Meyer and Kieras, 1997; Sigman and Dehaene, 2008; Watanabe and Funahashi, 2018). Consequently, interpretation of reduced dual-task performance is rather difficult which represents an important limitation of this study.

However, it appears that dorsolateral prefrontal cortex, a brain region responsible for higher-level task executive, is bringing several deficits together, since it is: 1) structurally degraded with aging), 2) likely responsible for reduced explicit adaptation (Anguera et al., 2011), 3) likely involved in dual-task execution (Leone et al., 2017; Watanabe and Funahashi, 2018), and 4) important for working memory capacity (Rypma and D’Esposito, 2000). This brain region appears to be an interesting target when designing future studies that investigate the cause for explicit strategy decline with aging.

## Conclusion and outlook

In this paper, we found that older adults did not have higher motor or cognitive dual-task costs during unperturbed reaching movements and that the observed dual-task costs could not explain the age-related decline in the explicit component of motor adaptation (Heuer and Hegele, 2008; Vandevoorde and Orban de Xivry, 2019). Rather, we observed that the explicit component of motor adaptation was reliably associated with working memory capacity. This suggests that the amount of working memory resources of an individual is a good predictor for the magnitude of the explicit component during a visuomotor rotation task. Our study leaves several questions unanswered: 1) Why do cognitive abilities contribute to learning in simple motor tasks but not in motor adaptation? Is this linked to the nature of the task itself? 2) How can we train people to overcome this decline? One possibility is to reduce working memory demands in older adults during motor learning (Rajeshkumar and Trewartha, 2019). Another possibility is to train working memory capacity in older adults (Anguera et al., 2012). 3) Are the same mechanisms leading to a decline in the explicit component and to working memory or is there a direct causal link between the decline in working memory and the decline in the explicit component of adaptation?

Working memory can be approached as a multicomponent (Baddeley and Hitch, 1974) or as a state-based model (Esposito and Postle, 2015). Therefore, to further explore the nature of the explicit strategy decline in older adults, it might be useful to relate it to different states or components of working memory. This approach would require the design of novel paradigms that can dissociate working memory components with respect to explicit strategy development, an approach recently initiated by McDougle and Taylor (2019). This seems a promising approach to gain further insights in the decline of explicit strategy with aging.

In order to overcome the decline of explicit strategy with aging, it needs to be tested whether targeted cognitive training might resolve some of the deficits. Another possibility is the application of non-invasive brain stimulation, which is a technique that appears to be suited to restore some of the age-related deficits (Hardwick and Celnik, 2014; Orban de Xivry and Shadmehr, 2014; Grimaldi et al., 2016). A two-fold approach is possible to restore the decline of explicit strategy in elderly: Either directly, by stimulation of a region such as dorsolateral prefrontal cortex that might temporary boost some working memory resources (Seidler et al., 2017), or indirectly, by stimulation of the cerebellum that might boost, the already intact, implicit component of motor adaptation. Finally, we expect that the observed age-related decline of explicit strategy is having widespread consequences on other features and components of motor adaptation such as reduced generalization, increased interference of motor memories, reduced savings and reduced reinforcement learning.

## Acknowledgements

This work was supported by an internal grant of the KU Leuven (STG/14/054) and by the FWO (1519916N). We thank Heleen Verbist for helping us with the data collection and Rob Hardwick for English language editing.

## References

Anguera J a, Reuter-Lorenz P a, Willingham DT, Seidler RD (2010) Contributions of spatial working memory to visuomotor learning. J Cogn Neurosci 22:1917–1930.

Anguera J a, Reuter-Lorenz P a, Willingham DT, Seidler RD (2011) Failure to engage spatial working memory contributes to age-related declines in visuomotor learning. J Cogn Neurosci 23:11–25.

Anguera JA, Bernard JA, Jaeggi SM, Buschkuehl M, Benson BL, Jennett S, Humfleet J, Reuter-lorenz PA, Jonides J, Seidler RD (2012) The effects of working memory resource depletion and training on sensorimotor adaptation. Behav Brain Res 228:107–115.

Benson BL, Anguera JA, Seidler RD (2011) A spatial explicit strategy reduces error but interferes with sensorimotor adaptation. J Neurophysiol 105:2843–2851.

Birren J, Fisher L (1995) Aging and Speed of Behavior: Possible Consequences for Psychological Functioning. Annu Rev Psychol 46:329–353.

Boisgontier MP, Beets IAM, Duysens J, Nieuwboer A, Krampe RT, Swinnen SP (2013) Age-related differences in attentional cost associated with postural dual tasks: Increased recruitment of generic cognitive resources in older adults. Neurosci Biobehav Rev 37:1824–1837.

Cappell KA, Gmeindl L, Reuter-Lorenz PA (2010) Age differences in prefontal recruitment during verbal working memory maintenance depend on memory load. Cortex 46:462–473.

Cashaback JGA, Mcgregor HR, Mohatarem A, Gribble L (2017) Dissociating error-based and reinforcement-based loss functions during sensorimotor learning.:1–28.

Christou AI, Miall RC, McNab F, Galea JM (2016) Individual differences in explicit and implicit visuomotor learning and working memory capacity. Sci Rep 6:36633.

Curtis CE, Esposito MD (2003) Persistent activity in the prefrontal cortex during working memory. 7:415–423.

Darling WG, Cooke JD, Brown SH (1989) Control of Simple Arm Movements in Elderly Humans. 10:149–157.

Davis SW, Kragel JE, Madden DJ, Cabeza R (2012) The architecture of cross-hemispheric communication in the aging brain: Linking behavior to functional and structural connectivity. Cereb Cortex 22:232–242.

Doumas M, Smolders C, Krampe RT (2008) Task prioritization in aging: Effects of sensory information on concurrent posture and memory performance. Exp Brain Res 187:275–281.

Ebersbach G, Dimitrijevic MR, Poewe W (2011) Influence of Concurrent Tasks on Gait: A Dual-Task Approach. Percept Mot Skills 81:107–113.

Eriksen BA, Eriksen CW (1974) Effects of noise letters upon the identification of a target letter in a nonsearch task. Percept Psychophys 16:143–149.

Esposito MD, Postle BR (2015) The Cognitive Neuroscience of Working Memory.

Folstein M, Folstein S, McHugh P (1975) A practical state method for. 12:189–198.

Fozard JL, Vercruyssen M, Reynolds SL, Hancock PA, Quilter RE (1994) Age differences and changes in reaction time: The Baltimore longitudinal study of aging. Journals Gerontol 49.

Galea JM, Sami SA, Albert NB, Miall RC (2010) Secondary tasks impair adaptation to step- and gradual-visual displacements.:473–484.

Gazzaley A, Cooney JW, Rissman J, Esposito MD (2005) Top-down suppression deficit underlies working memory impairment in normal aging. 8:1298–1301.

Goble DJ., Coxon JP., Wenderoth N, Van Impe A, Swinnen SP (2009) Proprioceptive sensibility in the elderly: Degeneration, functional consequences and plastic-adaptive processes. Neurosci Biobehav Rev 33:271–278.

Grady C (2012) The cognitive neuroscience of ageing. Nat Rev Neurosci 13:491–505.

Grimaldi G, Argyropoulos GP, Bastian A, Cortes M, Davis NJ, Edwards DJ, Ferrucci R, Fregni F, Galea JM, Hamada M, Manto M, Miall RC, Morales-Quezada L, Pope PA, Priori A, Rothwell J, Tomlinson SP, Celnik P (2016) Cerebellar Transcranial Direct Current Stimulation (ctDCS): A Novel Approach to Understanding Cerebellar Function in Health and Disease. Neuroscientist 22:83–97.

Guttmann CRG, Jolesz FA, Kikinis R, Killiany RJ, Moss MB, Sandor T, Albert MS (1998) White matter changes with normal aging. Neurology 50:972–978.

Hardwick RM, Celnik P a (2014) Cerebellar direct current stimulation enhances motor learning in older adults. Neurobiol Aging 35:1–5.

Heuer H, Hegele M (2008) Adaptation to visuomotor rotations in younger and older adults. Psychol Aging 23:190–202.

Heuninckx S, Wenderoth N, Debaere F, Peeters R, Swinnen SP (2005) Neural Basis of Aging: The Penetration of Cognition into Action Control. J Neurosci 25:6787–6796.

Heuninckx S, Wenderoth N, Swinnen SP (2008) Systems neuroplasticity in the aging brain: recruiting additional neural resources for successful motor performance in elderly persons. J Neurosci 28:91–99.

Huang J, Hegele M, Billino J (2018) Motivational modulation of age-related effects on reaching adaptation. Front Psychol 9:1–13.

Huxhold O, Li S, Schmiedek F, Lindenberger U (2006) Dual-tasking postural control : Aging and the effects of cognitive demand in conjunction with focus of attention. 69:294–305.

Jost K, Bryck RL, Vogel EK, Mayr U (2011) Are Old Adults Just Like Low Working Memory Young Adults ? Filtering Efficiency and Age Differences in Visual Working Memory.:1147–1154.

Jubrias SA, Odderson IR, Esselman PC, Conley KE (1997) Decline in isokinetic force with age: Muscle cross-sectional area and specific force. Pflugers Arch Eur J Physiol 434:246–253.

Karbach J, Verhaeghen P (2014) Making Working Memory Work : A Meta-Analysis of Executive-Control and Working Memory Training in Older Adults.

Keisler A, Shadmehr R (2010) A Shared Resource between Declarative Memory and Motor Memory. 30:14817–14823.

Ketcham CJ, Seidler RD, Gemmert AWA Van, Stelmach GE (2002) Age-Related Kinematic Differences as Influenced by Task Difficulty, Target Size, and Movement Amplitude. 57:54–64.

Klingberg T (2010) Training and plasticity of working memory. Trends Cogn Sci 14:317–324.

Kopp B, Mattler U, Rist F (1994) Selective attention and response competition in schizophrenic patients. Psychiatry Res 53:129–139.

Krakauer JW, Ghilardi M, Mentis M, Barnes A, Veytsman M, Eidelberg D, Ghez C, John W, Ghilardi M, Mentis M, Veytsman M, Eidelberg D, Ghez C (2004) Differential Cortical and Subcortical Activations in Learning Rotations and Gains for Reaching : A PET Study. 3:924–933.

Leone C, Feys P, Moumdjian L, D’Amico E, Zappia M, Patti F (2017) Cognitive-motor dual-task interference: A systematic review of neural correlates. Neurosci Biobehav Rev 75:348–360.

Li KZH, Lindenberger U (2002) Relations between aging sensory / sensorimotor and cognitive functions. 26:777–783.

Lindle RS, Metter EJ, Lynch NA, Fleg JL, Fozard JL, Tobin J, Roy TA, Hurley BF (1997) Age and gender comparisons of muscle strength in 654 women and men aged 20-93 yr. J Appl Physiol 83:1581–1587.

López-Otín C, Blasco MA, Partridge L, Serrano M, Kroemer G (2013) The hallmarks of aging. Cell 153:1194–1217.

Ma H-I, Hwang W-J, Lin K-C (2009) The effects of two different auditory stimuli on functional arm movement in persons with Parkinson’s disease : a dual-task paradigm.:229–237.

Madden DJ, Costello MC, Dennis NA, Davis SW, Shepler AM, Spaniol J, Bucur B, Cabeza R (2010) Adult age differences in functional connectivity during executive control. Neuroimage 52:643–657.

Malone LA, Bastian AJ (2010) Thinking about walking: effects of conscious correction versus distraction on locomotor adaptation. J Neurophysiol 103:1954–1962.

Martin JA, Buckwalter JA (2002) Aging, articular cartilage chondrocyte senescence and osteoarthritis. Biogerontology 3:257–264.

Mattay VS, Fera F, Tessitore A, Hariri AR, Berman KF, Das S, Meyer-Lindenberg A, Goldberg TE, Callicott JH, Weinberger DR (2006) Neurophysiological correlates of age-related changes in working memory capacity. Neurosci Lett 392:32–37.

Mazzoni P, Krakauer JW (2006) An implicit plan overrides an explicit strategy during visuomotor adaptation. J Neurosci 26:3642–3645.

McDougle SD, Bond KM, Taylor JA (2015) Explicit and Implicit Processes Constitute the Fast and Slow Processes of Sensorimotor Learning. J Neurosci 35:9568–9579.

Mcdougle SD, Ivry RB, Taylor JA (2016) Taking Aim at the Cognitive Side of Learning in Sensorimotor Adaptation Tasks. Trends Cogn Sci 20:535–544.

McDougle SD, Taylor JA (2019) Dissociable cognitive strategies for sensorimotor learning. Nat Commun.

McNab F, Klingberg T (2008) Prefrontal cortex and basal ganglia control access to working memory. Nat Neurosci 11:103–107.

Melzer I, Benjuya N, Kaplanski J (2001) Age-related changes of postural control: Effect of cognitive tasks. Gerontology 47:189–194.

Meyer DE, Kieras DE (1997) A Computational Theory of Executive Cognitive Processes and Multiple-Task Performance: Part 1. Basic Mechanisms. Psychol Rev 104:3–65.

Monsell S (2003) Task switching. 7:134–140.

Morehead JR, Qasim SE, Crossley MJ, Ivry R (2015) Savings upon Re-Aiming in Visuomotor Adaptation. J Neurosci 35:14386–14396.

Morehead JR, Taylor JA, Parvin D, Ivry RB (2017) Characteristics of Implicit Sensorimotor Adaptation Revealed by Task-irrelevant Clamped Feedback. J Cogn Neurosci 26:194–198.

Morrison AB, Chein JM (2011) Does working memory training work ? The promise and challenges of enhancing cognition by training working memory.:46–60.

Moscovitch M, Winocur G (1995) Frontal Lobes, Memory, and Aging.

Nagel IE, Preuschhof C, Li S-C, Nyberg L, Bäckman L, Lindenberger U, Heekeren HR (2011) Load modulation of BOLD response and connectivity predicts working memory performance in younger and older adults. J Cogn Neurosci 23:2030–2045.

Oldfield RC (1971) The assessment and analysis of handedness: The Edinburgh inventory. Neuropsychologia 9:97–113.

Orban de Xivry J-J, Shadmehr R (2014) Electrifying the motor engram: effects of tDCS on motor learning and control. Exp Brain Res 232:3379–3395.

Ota M, Obata T, Akine Y, Ito H, Ikehira H, Asada T, Suhara T (2006) Age-related degeneration of corpus callosum measured with diffusion tensor imaging. 31:1445–1452.

Owsley C (2011) Aging and vision. Vision Res 51:1610–1622.

Pratt J, Neggers B (2008) Inhibition of return in single and dual tasks : Examining saccadic, keypress, and pointing responses. 70:257–265.

Rajeshkumar L, Trewartha KM (2019) Advanced spatial knowledge of target location eliminates age-related differences in early sensorimotor learning. Exp Brain Res.

Raz N, Lindenberger U, Rodrigue KM, Kennedy KM, Head D, Williamson A, Dahle C, Gerstorf D, Acker JD (2005) Regional brain changes in aging healthy adults: General trends, individual differences and modifiers. Cereb Cortex 15:1676–1689.

Raz N, Rodrigue KM (2006) Differential aging of the brain : Patterns, cognitive correlates and modifiers. 30:730–748.

Reuter-Lorenz PA, Campbell KA (2008) Neurocognitive ageing and the Compensation Hypothesis. Curr Dir Psychol Sci 17:177–182.

Rypma B, D’Esposito M (2000) Isolating the neural mechanisms of age-related changes in human working memory. Nat Neurosci 3:509–515.

Salthouse TA (2000) Aging and measures of processing speed. Biol Psychol 54:35–54.

Schneider-Garces NJ et al. (2009) Span, CRUNCH, and Beyond: Working Memory Capacity and the Aging Brain. J Cogn Neurosci 22:655–669.

Seidler RD, Bernard JA, Burutolu TB, Fling BW, Gordon MT, Gwin JT, Kwak Y, Lipps DB (2010) Motor control and aging: Links to age-related brain structural, functional, and biochemical effects. Neurosci Biobehav Rev 34:721–733.

Seidler RD, Gluskin BS, Greeley B (2017) Right prefrontal cortex transcranial direct current stimulation enhances multi-day savings in sensorimotor adaptation. J Neurophysiol 117.

Serrien DJ, Swinnen SP, Stelmach GE (2000) Age-related deterioration of coordinated interlimb behavior. Journals Gerontol - Ser B Psychol Sci Soc Sci 55:P295–P303.

Shadmehr R, Holcomb HH (1997) Neural correlates of motor memory consolidation. Science 277:821–825.

Shadmehr R, Smith M a, Krakauer JW (2010) Error correction, sensory prediction, and adaptation in motor control. Annu Rev Neurosci 33:89–108.

Sigman M, Dehaene S (2008) Brain Mechanisms of Serial and Parallel Processing during Dual-Task Performance. J Neurosci 28:7585–7598.

Spellman T, Rigotti M, Ahmari SE, Fusi S, Gogos JA, Gordon JA (2015) Hippocampal-prefrontal input supports spatial encoding in working memory. Nature 522:309–314.

Taylor J a., Krakauer JW, Ivry RB (2014) Explicit and Implicit Contributions to Learning in a Sensorimotor Adaptation Task. J Neurosci 34:3023–3032.

Taylor JA, Thoroughman KA (2008) Motor Adaptation Scaled by the Difficulty of a Secondary Cognitive Task. 3.

Van Halewyck F, Lavrysen A, Levin O, Elliott D, Helsen WF (2015) The impact of age and physical activity level on manual aiming performance. J Aging Phys Act 23:169–179.

Vandevoorde K, Orban de Xivry J-J (2019) Internal model recalibration does not deteriorate with age while motor adaptation does. Neurobiol Aging 80:138–153.

Vogel EK, Mccollough AW, Machizawa MG (2005) Neural measures reveal individual differences in controlling access to working memory. Nature 438:500–503.

Watanabe K, Funahashi S (2018) Toward an understanding of the neural mechanisms underlying dual-task performance: Contribution of comparative approaches using animal models. Neurosci Biobehav Rev 84:12–28.

Werner S, Van Aken BC, Hulst T, Frens MA, Van Der Geest JN, Strüder HK, Donchin O (2015) Awareness of sensorimotor adaptation to visual rotations of different size. PLoS One 10:1–18.

Yan JH, Thomas JR, Stelmach GE (1998) Aging and Rapid Aiming Arm Movement Control Movement Control. 4657.

